# The Actin regulator Mena promotes Wnt signalosome endocytosis and Wnt signalling

**DOI:** 10.1101/2025.11.24.690281

**Authors:** Sheng-yuan Wu, Marcela M. Moreno, Anna Noble, Amirul Haziq Azwan, Matthew Guille, Karen J. Liu, Matthias Krause

## Abstract

Wnt signaling controls embryonic development and tissue maintenance. Endocytosis of Wnt-receptors is required for signalling, yet uptake mechanisms remain poorly understood. Here, we identify the actin regulator and Ena/VASP protein, Mena, as a key mediator. Upon Wnt stimulation, Mena redistributes from focal adhesions to signalosomes, Wnt-receptor clusters. Mena directly binds Wnt-coreceptors LRP5/6 in a phosphorylation-dependent manner increasing Wnt signal transduction. Sequestration of Ena/VASP proteins impedes *in vivo* Wnt activation driving *Xenopus* embryonic development. We resolve previous controversies by showing that Wnt3a triggers rapid Clathrin-Mediated and Fast Endophilin-Mediated LRP6 endocytosis at low concentrations sufficient for Wnt activation. This efficient endocytosis requires Ena/VASP proteins and is specifically promoted by Mena. Our results suggest Mena as a crucial mediator of Wnt signalosome endocytosis thus promoting canonical Wnt signalling.

## Main Text

Wnt-signalling orchestrates fundamental processes from embryonic development to tissue maintenance with aberrations driving cancers and developmental disorders. Canonical Wnt-signalling requires tight regulation of levels of the transcriptional co-activator β-catenin by the β-catenin-destruction complex (DC), which contains Axin, APC, GSK3 and CK1. Upon Wnt ligand binding to Frizzled receptors and Wnt co-receptors LRP5 and LRP6, the latter two undergo phosphorylation by serine/threonine kinases GSK3 and CK1, creating binding sites for recruitment of Axin and the associated DC (*1–5*). Their clustering through Disheveled (Dvl) leads to the formation of signalosomes, membrane-associated multimolecular complexes, which then undergo endocytosis (*6–11*). There is conflicting evidence about the role in endocytosis for Wnt signalling with some reports describing a positive role (*12–17*), some a negative role (*18–25*) and others no role for endocytosis (*26*) but instead proposing a role for macropinocytosis (*27*, *28*). One reason for the discrepancy may be that high concentrations of Wnt3a (50-200ng/ml purified Wnt3a) or Wnt3a conditioned media (containing approximately 200-400ng/ml) (*29*, *30*) have been used in these studies potentially inducing macropinocytosis non-specifically. However, low Wnt3a concentrations (10 ng/ml) are sufficient for β-catenin stabilization and Wnt activation (*29*, *30*). In addition, some studies used inhibitors which may be less specific than desired (*12*, *13*, *16*, *17*, *27*, *28*). Nevertheless, several studies using siRNA-mediated knockdown of key components of endocytic pathways or dominant negative dynamin (*13–17*, *19*, *31*), a key mediator of endocytosis (*32*), suggest a function for endocytosis in canonical Wnt signalling.

Thus, a combination of signalosome endocytosis (*10*, *11*, *13–17*, *19*, *31*) and direct interaction of GSK3 with phosphorylated LRP5/6 (*33*, *34*) may lead to inhibition of the DC allowing accumulation of enough nuclear β-catenin to drive Wnt target gene transcription (*1–5*). Therefore, a critical question remains: how are Wnt receptors endocytosed?

### The Ena/VASP proteins Mena and EVL promote WNT signalling

It is well established that upon Wnt stimulation, the Wnt co-receptors, LRP5 and LRP6 are phosphorylated by GSK3 and CK1 at five proline-rich motifs creating binding sites for Axin thereby causing DC recruitment to these receptors (*35*, *36*). We noticed that two sites in LRP5 and one in LRP6 overlap with a consensus motif, which in other proteins facilitate binding of the EVH1 domain of Ena/VASP proteins (Fig. 1A). Ena/VASP proteins (Mena, VASP, EVL) are actin filament elongators, important for cell migration (*37–39*). Ena/VASP proteins are implicated in ligand-induced clathrin-mediated uptake of a few specific receptors (Ephrin, EGF, and VEGF) (*40–42*) but a role in Wnt receptor endocytosis is unknown. Ena/VASP proteins harbour an N-terminal EVH1 domain mediating interactions with proline-rich ligands thereby ensuring recruitment to the leading edge of cells, focal adhesions, and sites of endocytosis. This is followed by its own proline-rich region and a C-terminal EVH2 domain which contains G- and F-actin binding sites that promote F-actin elongation as a processive actin polymerase, and a coiled-coil domain mediating tetramerization (*37–39*).

**Fig. 1.**
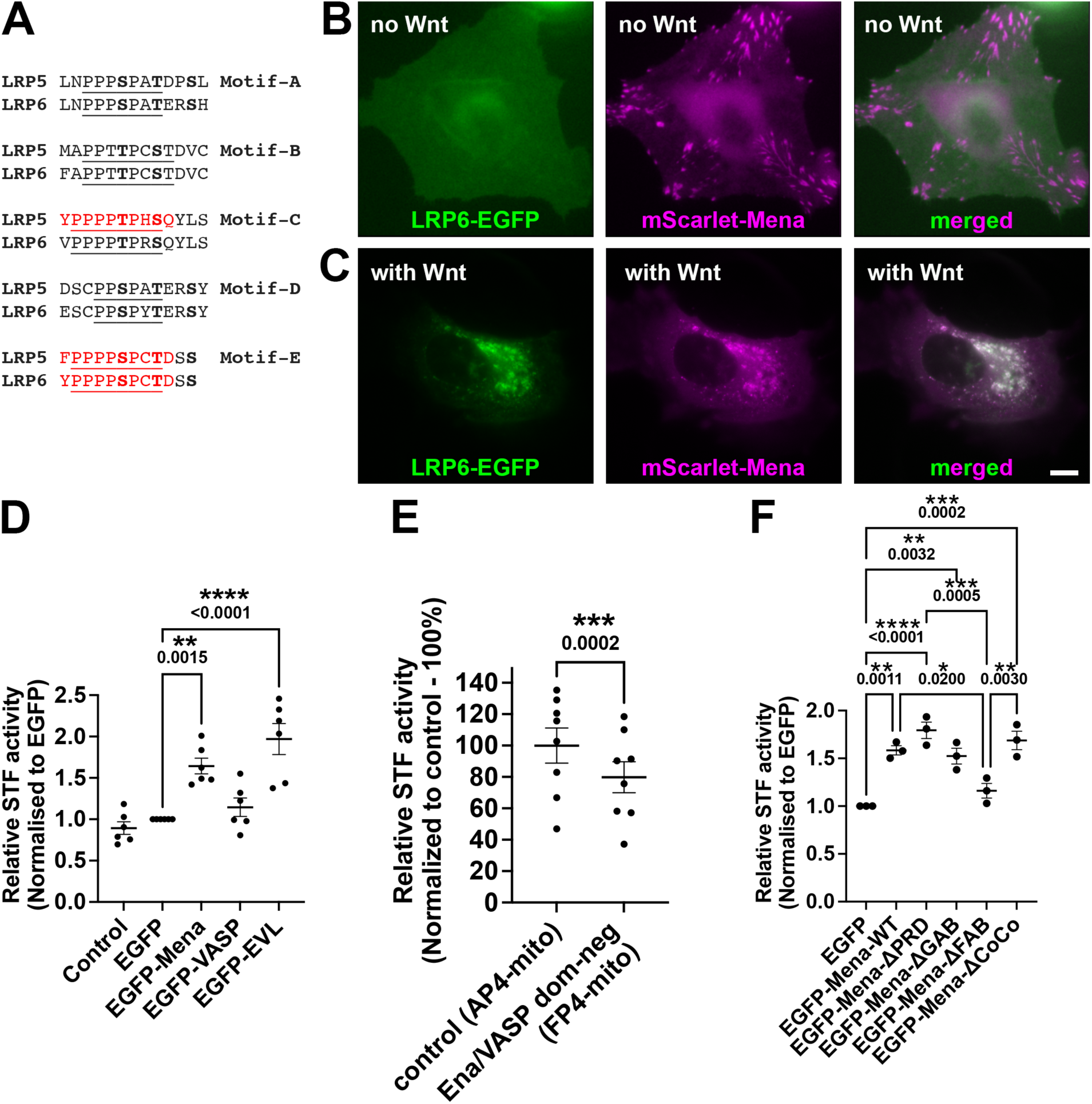
Mena re-localises to signalosomes and Mena and EVL promote WNT signalling. **(A)** Axin binding motifs A-E in LRP5 (NP_002326.2) and LRP6 (NP_001401173.1), single amino acid code, underlined letters indicate Axin binding sites; bold letters indicate serine or threonine phosphorylation by GSK3 or CK1; Ena/VASP EVH1 domain binding motifs: red amino acids. **(B,C)** HeLa cells transfected with LRP6-EGFP and mScarlet-Mena were imaged live without (B) and with (C) Wnt3a-conditioned media for 30 minutes. Scale bar = 10µm; three independent biological repeats. **(D-F)** Wnt activity was quantified using the STF assay in HEK293 cells stimulated with Wnt3a and transfected with **(D)** EGFP-tagged Mena, VASP, or EVL or EGFP only as negative control; **(E)** control AP4-mito or dominant-negative FP4-mito; **(F)** EGFP-tagged wild-type Mena or Mena mutants in the proline-rich (Mena-ΔPRD), G-actin binding site (Mena-ΔGAB), F-actin binding site (Mena-ΔFAB), with a deletion of the coiled-coil domain (Mena-ΔCoCo), or EGFP only as control. (D) One-way ANOVA, Dunnetts; (E) paired, two-tailed t-test; (F) One-way ANOVA, Tukey’s; 3-8 independent biological repeats.

Upon Wnt3a stimulation, Frizzled and LRP6 receptors together with the beta-catenin destruction complex form signalosomes, membrane-associated multimolecular complexes, which are then endocytosed (*6–9*). We examined a potential recruitment of Mena to signalosomes by transfecting LRP6-EGFP together with mScarlet-Mena into HeLa cells. As expected, we found that LRP6 resides diffusely in the plasma membrane and Mena localizes predominantly at focal adhesions (Fig. 1B). However, when we treated the cells for 30 minutes with Wnt3a conditioned media, we observed a dramatic re-localisation of Mena to LRP6-positive signalosomes (Fig. 1C).

To explore whether Ena/VASP proteins support Wnt transcriptional activation, we overexpressed individual Ena/VASP proteins in HEK293 cells together with either the canonical Wnt reporter SuperTopFLASH (STF), which contains β-catenin-dependent LEF/TCF binding sites driving luciferase, or the control where the LEF/TCF sites have been mutated (*49*). We observed that overexpression of Mena or EVL but not VASP caused about a two-fold increase in Wnt transcriptional activation (Fig. 1D).

Ena/VASP proteins normally localize to focal adhesions, the leading edge of the cell, and sites of endocytosis (*37–40*, *50*, *51*). To investigate a requirement of Ena/VASP proteins in Wnt activation we made use of a well-established knock-sideways strategy to delocalize Ena/VASP proteins away from their normal locations to mitochondria (*52*). Overexpression of this dominant-negative construct (FP4-mito) reduced Wnt-dependent transcriptional activation by 20% suggesting that Ena/VASP proteins support Wnt signalling (Fig. 1E). The reduction by 20% is similar to that for EGFR endocytosis where F-actin polymerisation contributes a similar percentage of the EGFR uptake (*40*). Ena/VASP requirement may be higher *in vivo* as F-actin polymerisation is essential for clathrin-mediated endocytosis on the apical side of epithelial cells due to higher membrane tension at the apical side (*53*). To explore whether Mena promotes Wnt signalling through its interaction with F-actin or through other interactors, we overexpressed Mena wild-type or Mena mutants carrying deletions in the proline-rich region, the coiled-coil domain or mutations in the G-actin or F-actin binding sites. Activity in Wnt signalling was then quantified using the STF assay. We observed that all Mena constructs led to an increase in Wnt activation, except Mena carrying mutations in the F-actin binding site (Fig. 1F). This indicates that Mena promotes Wnt signalling by increasing F-actin polymerization.

### A novel LRP6 - Ena/VASP interaction is increased by GSK3 and CK1 phosphorylation

Because Mena translocates to signalosomes upon Wnt stimulation and promotes Wnt signalling, we explored whether Mena interacts directly with the Wnt coreceptors LRP5/6 in an inducible manner. Known EVH1 binding sites in other proteins are specific proline-rich peptide motifs: F/Y/W/LPXΦPX(E/D)(E/D)EL (X = any and Φ = hydrophobic amino acid) (*43–46*) and the acidic amino acids, glutamic acid (E) and aspartic acid (D) can mimic phosphorylated serine and threonine residues (*47*). Therefore, we tested whether Ena/VASP proteins may bind to LRP5 and LRP6 in a phosphorylation-dependent manner. Ena/VASP proteins have a c-terminal coiled-coil domain mediating tetramerization and thus increase EVH1 binding through increased avidity. We thus fused the Mena EVH1 domain to its own c-terminal coiled-coil domain using a previously established glycine-rich linker (*46*) and verified the functionality of the coiled-coil interaction (fig. S1). We chose Mena to test whether its EVH1 domain interacts specifically with LRP5 or LRP6 because it is the evolutionarily most conserved family member being the sole orthologue in invertebrates (*48*). We *in vitro* phosphorylated purified, bead-immobilised GST-LRP5 or -LRP6 with purified GSK3, CK1 or both and verified their phosphorylation (Fig. 2A,B; fig. S1). Incubation of these GST-LRP6 beads with purified Strep-tag-Mena-EVH1-CC protein revealed that dual phosphorylation of LRP6 by GSK3 and CK1 increases its interaction with the Mena-EVH1-CC twenty-fold (Fig. 2A,B). A similar GSK3/CK1 phosphorylation-induced increase has been previously reported for the LRP6-destruction complex interaction (*35*, *36*). We also observed an increase in interaction with Mena-EVH1-CC upon phosphorylation of LRP5 albeit to a moderate degree (fig. S1).

**Fig. 2.**
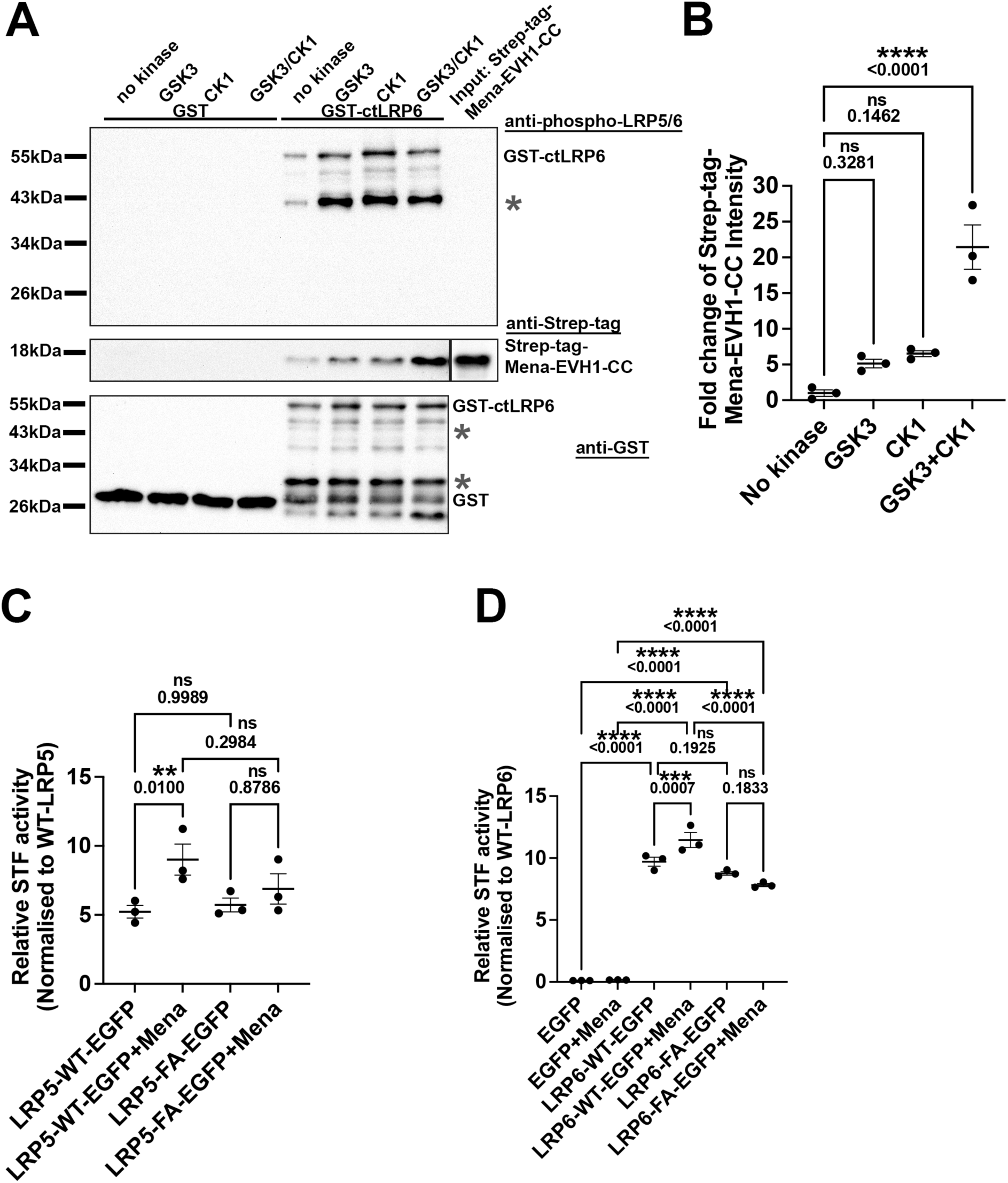
The LRP6 - Mena interaction is increased by GSK3 and CK1 phosphorylation and Mena promotes WNT signalling through an interaction with LRP6. **(A)** Immobilised, purified GST-tagged intracellular c-terminus of LRP6 or GST only as negative control was *in vitro* phosphorylated with either GSK3, CK1, or both, and incubated with purified Mena Strep-tag-EVH1-CC. Phosphorylation was assessed using an anti-phospho-LRP6 antibody (top panel). Interaction was visualized in a western blot against the strep-tag (middle panel) or GST (lower panel). Note some degradation of GST-ctLRP6 occurs due to its unstructured nature indicated in (B) as *. **(B)** Quantification of Strep-tag-Mena-EVH1-CC band intensity from (A). One-way ANOVA, Tukey’s; Mean ± SEM; three independent biological repeats. **(C,D)** Wnt activity was quantified using the STF assay in LRP5/6 double knockout HEK293 cells stimulated with Wnt3a and transfected with EGFP only as control or LRP5- or LRP6-EGFP tagged wild-type or Ena/VASP-binding deficient LRP5 or LRP6 (LRP5-FA-EGFP or LRP6-FA-EGFP) with and without Myc-tagged Mena. (C,D) One-way ANOVA, Tukey’s; 3 independent biological repeats.

### Mena promotes WNT signalling through an interaction with LRP6

To test whether Ena/VASP proteins increase canonical Wnt signalling through an interaction with LRP5 or LRP6, we generated HEK293 cells carrying LRP5/6 mutants lacking EVH1 binding. To do this, we first generated LRP5/6 double knockout HEK293 cell lines using CRISPR-Cas9 (validation, fig. S2-S3). We then restored LRP5/6 by introducing EGFP-tagged wild-type or versions of LRP5 or LRP6 where the EVH1 binding sites are mutated but retain an intact Axin interacting motif. This was done by mutating tyrosine or phenylalanine in the LRP5 motifs C and E or the tyrosine in the LRP6 motif E to alanine (Fig. 1A). Mutating the F/Y/W/L in the EVH1 binding F/Y/W/LPXΦPX(E/D)(E/D)EL recognition site to alanine has previously been shown to cause loss of interaction (*43*, *44*). We confirmed that re-expressed wild-type or mutated LRP6 was seen at near endogenous levels in LRP5/6 double knockout HEK293 cells (fig. S3). GFP-only was used as a control.

We then overexpressed Myc-tagged Mena with or without concomitant Wnt3a stimulation and measured Wnt transcriptional activation in a STF assay. We found that, as expected, in the absence of LRP5/6, Wnt3a stimulation did not increase Wnt reporter activation (Fig. 2D). Also, as expected from the above experiments, we found that Mena overexpression increased Wnt reporter activation when we re-expressed wild-type LRP5 or LRP6 with Wnt3a stimulation. However, the Mena-dependent Wnt activation was not seen in cells expressing LRP5 or LRP6 lacking the Ena/VASP binding sites (LRP5-FA-EGFP or LRP6-FA-EGFP), demonstrating the requirement for LRP5/6 interaction (Fig. 2C,D).

### LRP6 endocytosis is mediated at low Wnt3a stimulation by CME and FEME

To investigate the mechanism of Ena/VASP function in canonical Wnt signalling we explored a role in endocytosis as Mena translocates to signalosomes and is implicated in endocytosis of other receptors (*40–42*). However, there is conflicting evidence about the role for endocytosis in Wnt signalling (*12–28*). One reason for the discrepancy may be that some experiments used high concentrations of Wnt3a (50-400ng/ml Wnt3a) (*29*, *30*) despite lower Wnt3a concentrations (10 ng/ml) being sufficient for β-catenin stabilization and Wnt activation (*29*, *30*).

Therefore, we first explored the concentration- and timing-dependence of LRP6 endocytosis. To quantify the amount of endogenous LRP6 endocytosed by cells upon Wnt3a stimulation, we set up an endocytosis assay using hTERT-RPE1 cells which have an intact canonical Wnt pathway (*54*). We biotinylated all surface proteins while keeping the cells on ice to prevent endocytosis. Cells were then stimulated with Wnt3a conditioned medium for for 2, 5, 10, or 20 minutes at 37°C followed by removal of the biotin from the cell surface but not internalized proteins. After lysis, we then captured all endogenous LRP6 from the cell lysates on anti-LRP6-coated ELISA plates quantifying only the internalized, biotinylated LRP6 proteins, being careful to remain in the linear range (fig. S4). We found that already by 5 minutes the highest level of LRP6 endocytosis had occurred (Fig. 3A, fig. S4).

**Fig. 3:**
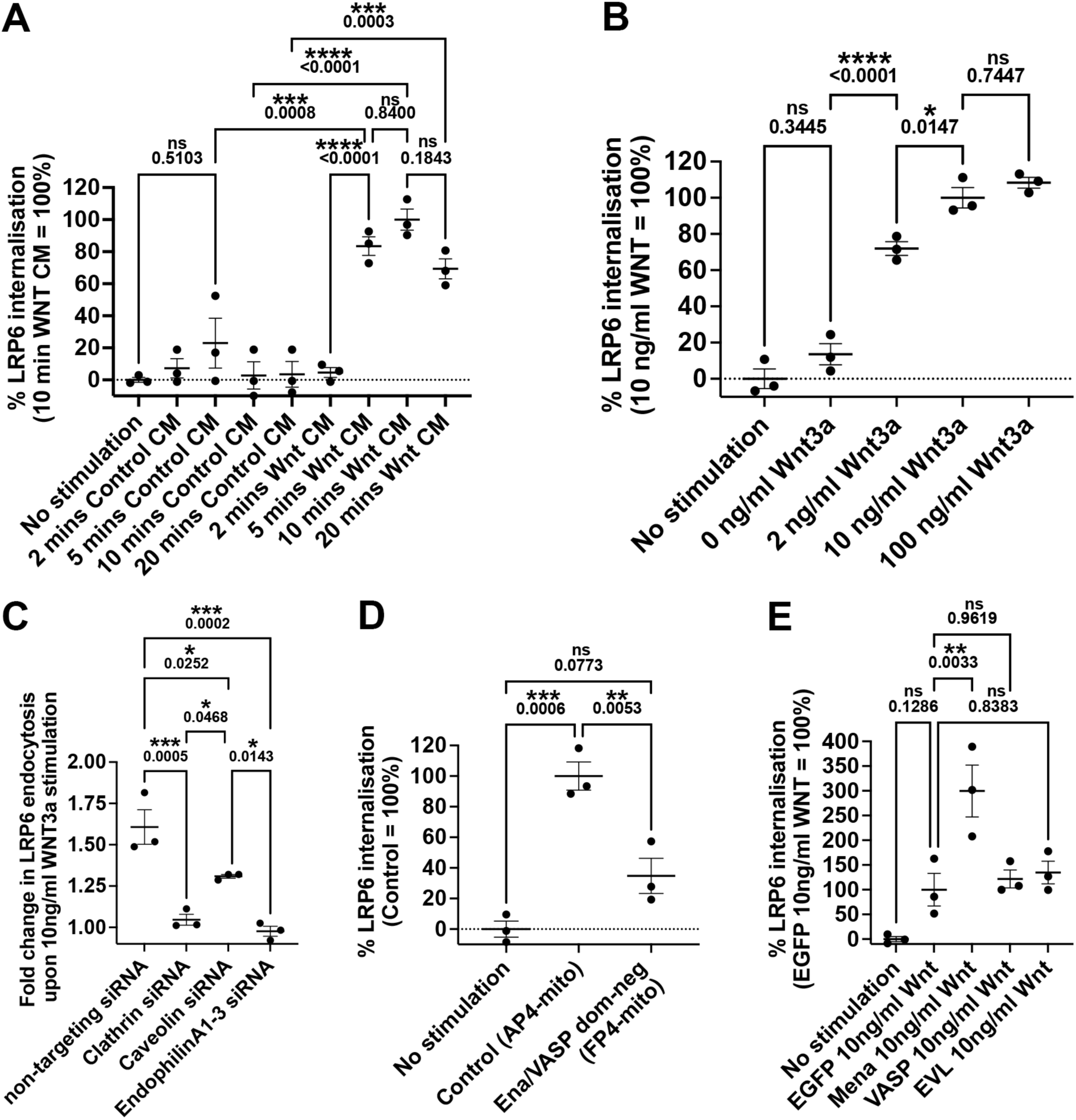
Mena promotes LRP6 endocytosis upon Wnt stimulation. **(A-E)** Endocytosis assay to quantify the uptake of endogenous LRP6 from hTERT-RPE1 cells. (A) Cells were stimulated for indicated times with control or Wnt3a conditioned media (CM). (B) Cells were stimulated for 5 minutes with the indicated concentrations of purified Wnt3a. (C) Cells were transfected with indicated siRNAs and then stimulated with 10 ng/ml Wnt3a for 5 minutes. (D,E) Cells were transfected with (D) control (AP4-mito) or Ena/VASP dominant negative (FP4-mito) plasmids or (E) EGFP-tagged Mena, VASP, EVL or EGFP only as negative control and stimulated with 10 ng/ml Wnt3a for 5 minutes. One-way ANOVA, (A-D) Tukey’s (E) Dunnetts; three independent biological repeats.

To evaluate whether lower concentrations of Wnt are sufficient for inducing LRP6 endocytosis we incubated cells with 2, 10 or 100 ng/ml purified Wnt3a for 5 minutes and found that as little as 2 ng/ml Wnt3a is sufficient to induce LRP6 endocytosis. This was further increased with 10 ng/ml Wnt3a, which appeared to achieve maximal stimulation since no further increase was seen with 100 ng/ml (Fig. 3B, fig. S4).

To explore the endocytosis pathway mediating LRP6 endocytosis at the established non-excess-Wnt conditions (10 ng/ml Wnt3a for 5 minutes), we knocked down clathrin heavy chain, caveolin-1, or all three endophilins (A1-3), which are required for Fast Endophilin-Mediated endocytosis (FEME) (knockdown validation, fig. S5). We found that Wnt3a induced LRP6 endocytosis was clathrin-mediated, partially dependent on caveolae and surprisingly also dependent on FEME (Fig. 3C, fig. S4). This suggests that Wnt-receptor endocytosis depends on at least two parallel pathways, CME and FEME, which may allow differentially signalling from distinct endosomal compartments.

We then asked whether Ena/VASP proteins contribute to LRP6 endocytosis. We again used the dominant-negative strategy to delocalise all Ena/VASP proteins to mitochondria and observed that endocytosis was severely reduced, indicating that Ena/VASP proteins are required for efficient LRP6 endocytosis (Fig. 3D, fig. S4). When we tested whether overexpression of Ena/VASP proteins affects LRP6 uptake, we found that only Mena causes a three-fold increase in LRP6 endocytosis (Fig. 3E, fig. S4). This suggests that Mena may activate Wnt signalling by increasing LRP6 endocytosis while EVL may be using a distinct mechanism.

### Ena/VASP proteins promote WNT signalling *in vivo*

To determine whether Ena/VASP proteins had similar roles in a developmental context *in vivo*, we turned to the *Xenopus* embryo, which has traditionally been used to evaluate effects on Wnt-driven development *in vivo*. Here, we used the Dorso-Anterior Index (DAI) (*55*) to quantify the extent of Wnt activation. As expected, ectopic introduction of Wnt8D mRNA into the animal hemisphere at the two-cell stage caused the majority of embryos to be strongly dorsalised (Fig 4C). Overexpressing of FP4-mito alone, or the control AP4-mito did not affect dorsal or anterior development of embryos. Consistent with the role for Ena/VASP in promoting Wnt signalling, co-expression of the dominant-negative FP4-mito construct was sufficient to counteract the severity of Wnt overexpression, leading to most embryos having clear axis elongation and restoration of eye and cement gland structures (Fig. 4A-E).

**Fig. 4:**
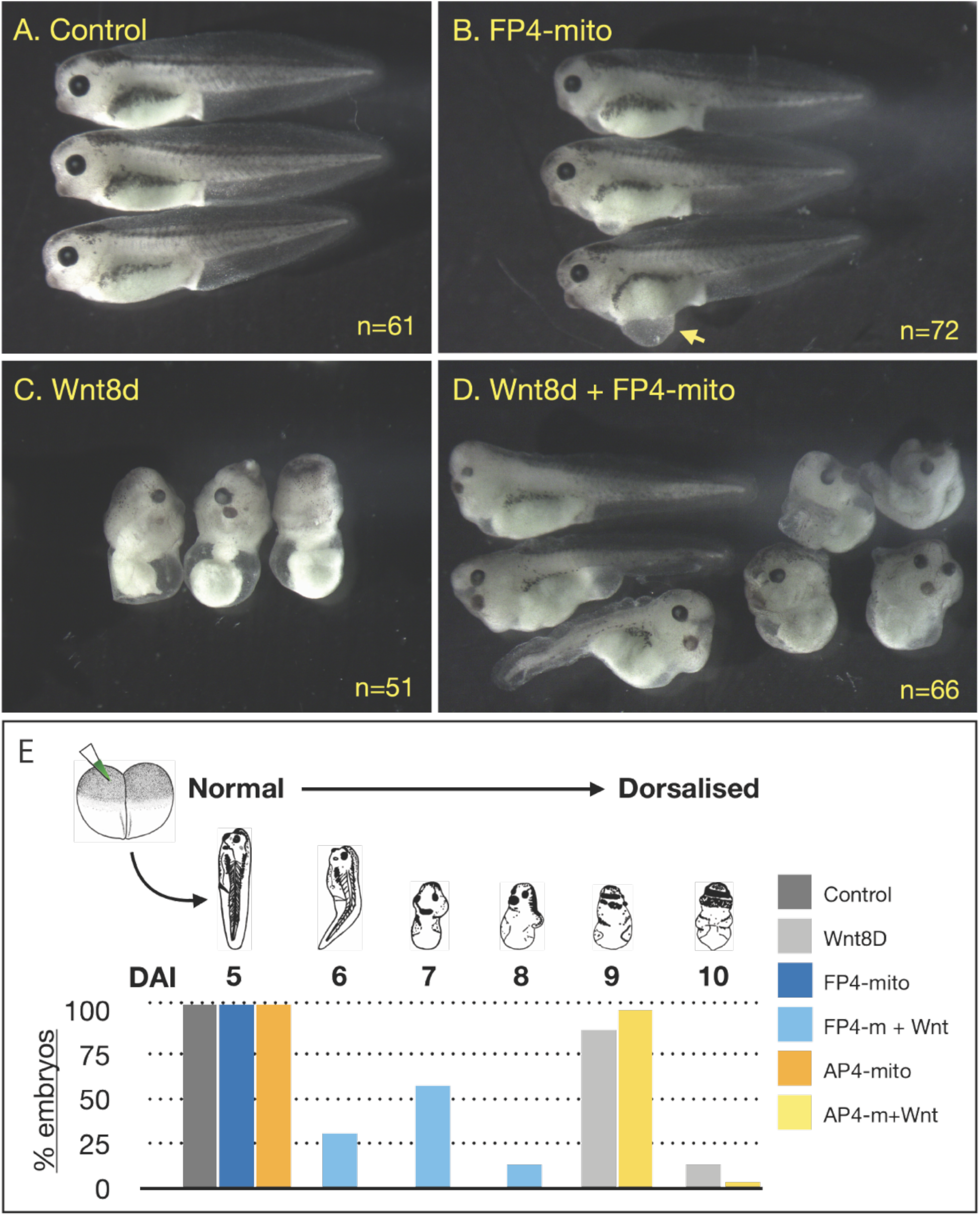
Overexpression of Wnt8 at Xenopus at 2-cell stage induces dorsalisation, which is counteracted by dominant-negative Ena/VASP (FP4-mito). **(A)** Control tadpoles. **(B)** Tadpoles injected at 2-cell stage with 1ng FP4-mito mRNA, note that FP4-mito alone causes some epithelial blistering, yellow arrow; (C) injected at 2-cell stage with 200pg Wnt8D; or (D) co-injected with 200pg Wnt8D and 1ng FP4 mRNA. **(E)** Distribution of phenotypes seen in tadpoles, based on Dorso-Anterior Index (DAI) after Kao and Elinson (*55*). AP4-mito control injections do not rescue dorsalisation caused by Wnt8D overexpression. N=51-72 embryos. Representative result of 4 independent experiments.

## Discussion

We discovered that Mena directly binds LRP6 Wnt co-receptors through a novel phosphorylation-dependent mechanism. The Mena EVH1 domain binds phosphorylated PPPSPxS motifs in LRP6, with dual GSK3/CK1 phosphorylation increasing LRP6 binding 20-fold, similar to LRP6-Axin regulation (*35*, *36*). This novel phosphorylation-controlled EVH1 domain binding contrasts with constitutive Ena/VASP interactions with other receptors (*56–59*).

Our functional studies revealed that only Mena and EVL, not VASP, enhance Wnt transcriptional activation, and this enhancement requires direct LRP5/6 binding as demonstrated using binding-deficient receptor mutants. It has been shown previously that individual Ena/VASP proteins can function independently of each other (*40*, *42*, *60–62*) as Mena or EVL only display weak hetero-tetramerization through their coiled-coil domains and mostly form independent Mena or EVL tetramers (*63*). We observed that Mena’s function depends solely on F-actin binding but not on its G-actin binding site nor its tetramerization, which is in agreement with that the F-actin binding site is the most important site for Mena function (*64–68*) indicating it acts here as an actin polymerase rather than through receptor clustering.

Mena binds only two motifs in LRP5 and one in LRP6, while Axin interacts with all five motifs (*69*). Recent identification of the membrane-proximal motif-A (Fig. 1A) as the primary Axin/GSK3 site (*34*) and Mena occupying distal motifs (Fig. 1A: motif-C and E), may explain why deletion of the distal motifs reduces Wnt signaling (*36*, *70*) suggesting that Mena and Axin cooperate rather than compete for LRP5/6 binding.

Our endocytosis studies resolve a longstanding controversy by showing that low Wnt3a concentrations sufficient for Wnt activation induce rapid LRP6 internalization via simultaneously engaging both CME and FEME. Previous conflicting reports used high Wnt3a concentrations, overexpressed receptors, or non-specific inhibitors, leading to conclusions favouring clathrin-mediated (*13*, *14*, *19–25*), caveolin-mediated (*16*, *17*, *19*, *31*), macropinocytic (*28*), or endocytosis-independent mechanisms (*26*). The dual CME/FEME pathway mirrors EGFR endocytosis (*71*) and may ensure robust endosomal signaling.

Ena/VASP proteins proved essential for efficient LRP6 endocytosis and Mena overexpression tripling uptake. We observed strong clathrin- and endophilin-dependent endocytosis with weaker caveolar involvement likely reflecting indirect membrane tension effects rather than direct caveolar endocytosis (*72–74*).

Significantly, Mena is a Wnt/β-catenin target gene (*75*), establishing a positive feedback loop where Wnt signaling upregulates Mena expression, which enhances LRP6 endocytosis and amplifies the signal. This mechanism provides a framework for understanding how cells achieve rapid sensitive responses to low-level Wnt signals through coordinated receptor phosphorylation, cytoskeletal recruitment, and dual-pathway endocytosis.

## Acknowledgments

We are grateful to Lisa Dobson, Wills Barrell, Jade Desjardins and other members of the Liu and Krause labs for valuable discussion and support. We are thanking the following for the kind gift of reagents: Emmanuel Boucrot, UCL, London, UK; Frank Gertler, MIT, Cambridge, MA, USA; Xi He, Harvard, Cambridge, MA, USA; Vivian Li; Francis Crick Institute, London, UK; David Virshup, Duke University, Durham, NC, USA; Randall Moon, University of Washington, Seattle, WA, USA. We thank Brian Stramer, Jeremy Green, Jody Rosenblatt, and Peter Bieling (King’s College London, UK) for critically reading the manuscript.

## Author contributions

Conceptualization: MK.

Methodology: MK, SW, KJL

Investigation: SW, MM, AN, AHA, KJL

Visualization: SW, KJL, MK

Funding acquisition: SW, MK, KJL, MG

Project administration: MK, KJL

Supervision: MK, KJL, MG

Writing – original draft: MK

Writing – review & editing: SW, MM, AN, MG, KJL, MK

## Data and materials availability

All data are available in the main text or the supplementary materials. Further information and requests relating to *Xenopus* resources and reagents, should be directed to KJL, and information and requests relating to mammalian experiments and biochemical resources and reagents should be directed to MK.

## Supplementary Materials

Materials and Methods

Figs. S1 to S5

Tables S1 to S2

References (*76–81*)

**The PDF file includes:**

Materials and Methods

Supplementary Text

Figs. S1 to S5

Tables S1 to S2

References 76-81

## Materials and Methods

### Molecular biology, plasmids and reagents

M50 Super 8x TOPFlash was a gift from Randall Moon (Addgene plasmid # 12456; http://n2t.net/addgene:12456; RRID: Addgene_12456). M51 Super 8x FOPFlash (TOPFlash mutant) was a gift from Randall Moon (Addgene plasmid # 12457; http://n2t.net/addgene:12457 ; RRID:Addgene_12457). pTwinStrep-DEST: A bacterial expression Gateway® (Invitrogen) destination vector with TwinStrep tag (pTwinStrep-DEST) was generated using pGEX-6P-1 (Cytiva) as the backbone and assembled using NEB HIFI with a synthetic DNA fragment (gBlocks^TM^, IDT) containing TwinStrep tag and overlapping sequences and the destination cassette from pDEST27 (Invitrogen) . The cDNA of dominant active mutant of GSK3 (pGSK-3 S9A) was cloned into pENTR3C (Invitrogen) by standard restriction enzyme-based cloning. The V405 4HA-CKIe (*76*) (a kind gift from David Virshup (Addgene plasmid # 13724 ; http://n2t.net/addgene:13724 ; RRID:Addgene_13724)) was used to clone the kinase domain of CK1e into pENTR3C (Invitrogen). The pENTR3C-Mena-EVH1-CC was generated using HiFi assembly (New England Biolabs) of Mena-EVH1, amplified from murine Mena cDNA (a kind gift from Frank Gertler, MIT, Cambridge, MA, USA), a synthetic DNA fragment (gBlocksTM, IDT) containing linker and coil-coiled domain sequence and pENTR3C (Invitrogen) as backbone. The c-termini of LRP5 or LRP6 were amplified from full-length LRP5 and LRP6 cDNAs (a kind gift from Xi He, Harvard, Cambridge, MA, USA) into pENTR3C (Invitrogen). Constructs for TwinStrep tagged GSK3, CK1 and Mena-EVH1-CC and GST tagged c-terminal LRP5 and LRP6 were generated by transferring cDNAs from pENTR3C into pTwinStrep-DEST or pDEST15 (Invitrogen) using Gateway® recombination (Invitrogen). Mena wild-type, Mena-ΔPRD. Mena-ΔGAP, Mena-ΔFAB, Mena-ΔCoCo in pMSCV-EGFP (*77*) were a kind gift from Frank Gertler, MIT, Cambridge, MA, USA.

FP4-mito, AP4-mito, murine VASP, Mena and EVL cDNAs (kind gifts from Frank Gertler, MIT, Cambridge, MA, USA) were cloned into pENTR3C (Invitrogen) and transferred into EGFP-tagged and mScarlet-I-tagged mammalian expression vectors using Gateway recombination. To mutate the EVH1 binding sites, the intracellular domains of LRP5 and LRP6 were replaced by synthetic DNA fragments (gBlocks^TM^, IDT) containing desired point mutations using NEB HiFi assembly (New England Biolabs).

### Antibodies

Primary antibodies. Mouse monoclonal anti-EGFP (Roche, 11814460001), WB (1:2000). Mouse monoclonal anti-Myc (Sigma Aldrich, M4439), WB (1:5000). Rabbit polyclonal anti-GST (Cell Signalling 2622), WB (1:1000). Mouse monoclonal anti-HSC70 (Santa Cruz SC7298), WB (1:1000). Rabbit monoclonal anti-LRP5 (Cell signalling 5731), WB (1:1000). Rabbit monoclonal anti-LRP6 (Cell signalling 3395), WB (1:1000). Rabbit monoclonal anti-Phospho-LRP6 (Cell signalling 2568), WB (1:1000). Mouse monoclonal Strep-MAB-Classic (IBA Lifescience 2-1507-001), WB (1:1000)

Secondary antibodies. HRP-goat anti-mouse IgG (Agilent, P0447), WB (1:2000). HRP-goat anti-rabbit IgG (Agilent, P0448), WB (1:2000). Anti-mouse IgG, HRP-linked Antibody (Cell signalling 7076), WB (1:1000). Anti-rabbit IgG, HRP-linked Antibody (Cell signalling 7074), WB WB (1:1000).

### Tissue culture

Human embryonic kidney 293 (HEK293FT) cells (Thermo Fisher Scientific, R70007) and the knockout cell lines derived from HEK293FT cells were culture in high-glucose Dulbecco’s modified Eagle’s medium (DMEM; Merck D6429) supplemented with 10% of fetal bovine serum (FBS, Gibco), 2mM L-glutamine (Merck, G7513), 100 unit/ml penicillin and 100µg/ml streptomycin (Merck; P0781) at 37°C in a 10% CO_2_ humidified incubator.

HeLa cells (ATCC CCL-2) were cultured in MEM supplemented with 1 x Non-essential Amino Acids (NEAA; Merck, M7145), 10% of FBS (Gibco), 2mM L-glutamine (Merck, G7513), 100 unit/ml penicillin and 100µg/ml streptomycin (Merck; P078) at 37°C in 5% CO_2_ humidified incubator.

hTERT-RPE1 cells (a kind gift of Emmanuel Boucrot, UCL; ATCC CRL-4000) were cultured in DMEM-F12Hams (Gibco, 11320022) supplemented with 10% of FBS (Gibco), 2mM L-glutamine (Merck, G7513), 100 unit/ml penicillin and 100µg/ml streptomycin (Merck; P078) at 37°C in 5% CO_2_ humidified incubator.

L-Wnt3a cells and L-control cells (a kind gift of Vivian Li; Francis Crick Institute) were cultured in DMEM supplemented with 125µg/ml Zeocin (ThermoFisher, J67140.XF), 10% of FBS (Gibco), 2mM L-glutamine (Merck, G7513), 100 unit/ml penicillin and 100µg/ml streptomycin (Merck; P078) at 37°C in 10% CO_2_ humidified incubator.

### Wnt3a conditioned medium

To prepare Wnt3a conditioned medium, L-Wnt3a cells were cultured in medium with Zeocin in a T175 flask. Once confluent, cells were split into 5x T175 flask without adding Zeocin. When the 5 flasks in medium without Zeocin are confluent (after 3-4 days). The cells were trypsinized and diluted into 600ml medium without Zeocin and plated onto 30 x 15cm tissue culture dishes (20ml/dish). After incubation at 37°C and 10% CO_2_ for one week, the medium was harvested and centrifuged at 300 rcf for 5 mins to remove floating cells. The Wnt3a conditioned medium were aliquoted into 50ml tubes and stored at 4°C. The activity of the conditioned medium was tested using the STF assay with the Dual-Glo® Luciferase Assay System (Promega).

### CRISPR-Cas9 KO

The sgRNAs targeting human *LRP5* and *LRP6* were designed to introduce premature stop codons within early exons. sgRNAs selection was performed using the CHOPCHOP web tool (https://chopchop.cbu.uib.no), prioritizing candidates with high predicted on-target activity and minimal off-target scores. Two sgRNAs were selected for each gene. The eSpCas9-plus nuclease (*78*) was used to mediate genome editing.

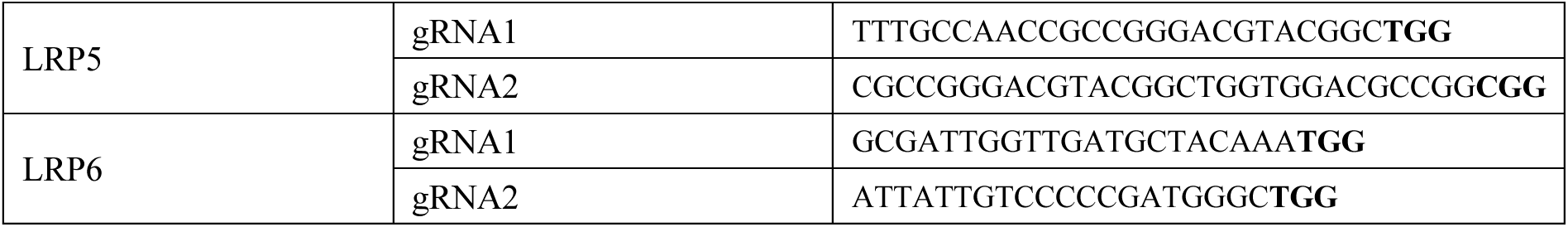

Forward and reverse oligonucleotides for each sgRNA were designed with BbsI-compatible overhangs (forward: 5’-caccgG-[sgRNA]-3’; reverse: 5’-aaac-[reverse complement sgRNA]-C-3’), annealed, and ligated into the BbsI sites of the peSpCas9-plus-T2A-Puro vector (*79*). Ligation reactions were treated with PlasmidSafe ATP-dependent DNase to remove unwanted products before transformation into E. coli. Positive clones were verified by Sanger sequencing and expanded by midi-prep purification.

HEK293FT cells were transfected with sgRNA-eSpCas9-plus-T2A-Puro constructs using Lipofectamine 2000. 48 hours after transfection, cells were selected with puromycin (4 µg/ml) for 48 h to enrich for transfected populations. To obtain clonal knockout lines, puromycin-resistant cells were subjected to limiting dilution.

To confirm gene knockout, genomic DNA was extracted from individual clones using the QIAamp DNA Mini Kit (Qiagen) according to the manufacturer’s instructions. The genomic region flanking the CRISPR–Cas9 target site was amplified by PCR, and the resulting products were subjected to Sanger sequencing. Sequencing chromatograms from each clone were compared with those from the wild-type control (HEK293FT) using the DECODR web tool (https://decodr.org/). Whole-cell lysates were prepared from expanded clones and analyzed by Western blot to assess *LRP5* and *LRP6* protein expression.

### Transfection and siRNA knockdown

A total of 1 × 10⁶ HeLa, HEK293FT, hTERT-RPE1, or HEK293FT-derived knockout cells were seeded in 6-well tissue culture plates in complete growth medium lacking penicillin–streptomycin. After 24 h incubation at 37 °C with appropriate CO₂ levels, plasmid transfection was performed using 2mg/ml polyethyleneimine (PEI) or Lipofectamine 2000 (for HEK293FT and hTERT-RPE1) or X-tremeGENE 360 (for HeLa).

For PEI transfection, 4 µg of plasmid DNA was diluted in 100 µl Opti-MEM in one microcentrifuge tube and 8 µl PEI was diluted in 100 µl Opti-MEM in a separate tube. Both mixtures were incubated at room temperature for 5 min, after which the PEI solution was added to the DNA solution and gently mixed. The DNA–PEI complexes were incubated for 20 min at room temperature before being added dropwise to the cells.

For Lipofectamine 2000 transfection, 4 µg of plasmid DNA was diluted in 125 µl Opti-MEM in one microcentrifuge tube and 10 µl Lipofectamine 2000 was diluted in 125 µl Opti-MEM in a separate tube. Both mixtures were incubated at room temperature for 5 min, after which the Lipofectamine solution was added to the DNA solution and gently mixed. The DNA and Lipofectamine mixture was incubated for 20 min at room temperature before being added dropwise to the cells.

For X-tremeGENE 360 transfection, 2 µg of plasmid DNA was diluted in 100 µl Opti-MEM in one microcentrifuge tube and 4 µl X-tremeGENE 360 was diluted in 100 µl Opti-MEM in a separate tube. Both mixtures were incubated at room temperature for 5 min, after which the X-tremeGENE 360 solution was added to the DNA solution and gently mixed. The DNA–X-tremeGENE 360 complexes were incubated for 20 min at room temperature before being added dropwise to the cells.

For Lipofectamine RNAiMAX transfection, 30 pmol of siRNA was diluted in 150 µl Opti-MEM in one microcentrifuge tube and 9 µl Lipofectamine RNAiMAX was diluted in 150 µl Opti-MEM in a separate tube. Both mixtures were incubated at room temperature for 5 min, after which the Lipofectamine RNAiMAX solution was added to the siRNA solution and gently mixed. The siRNA–Lipofectamine RNAiMAX complexes were incubated for 10 min at room temperature before being added dropwise to the cells. The siRNA oligos were transfected twice (reverse transfection at Day 1 and forward transfection at Day 2). The following siRNA oligos were used: clathrin heavy chain CLTC (Invitrogen Stealth, HSS174637, 5’-GAG UGC UUU GGA GCU UGU CUG UUU A-3’), caveolin 1 CAV1 (Invitrogen Stealth, HSS141467, 5’-CCC ACU CUU UGA AGC UGU UGG GAA A-3’), Endophilin A1 (SH3GL2) (Invitrogen Stealth, HSS109708, 5’-GGA UGA AGA GCU UCG UCA AGC UCU A-3’), Endophilin A2 (SH3GL1) (Invitrogen Stealth, HSS109707, 5’-CCC AAG AUC GCA GCU UCA UCG UCU U-3’) and Endophilin A3 (SH3GL3) (Invitrogen Stealth, HSS109712, 5’-CAA UGG AGU UUC CAC CAC CUC UGU A-3’). Transfected cells were incubated for 24–48 h prior to the following experiment.

### Pulldown, immunoprecipitation, and western blot

Cells samples were lysed in glutathione S-transferase (GST) buffer (50 mM Tris-HCL, pH 7.4, 200 mM NaCl, 1% NP-40, 2 mM MgCl2, 10% glycerol, 10mM NaF, 1mM Na3VO4 and EDTA-free protease inhibitor cocktail tablets (Roche)). Lysates were incubated on ice for 15 mins and centrifuged for 10 mins (17,000 xg at 4°C) to remove cellular debris. Protein concentration was measured using Pierce BCA protein assay kit following manufacture’s protocol (ThermoFisher Scientific). Absorbances were measured using a POLARstar Omega Plate Reader (SciQuip).

For pulldowns, glutathione-Sepharose beads (GE Healthcare), GFP-trap beads (Chromotek), or GFP-selector beads (NanoTag) were blocked with 1% BSA for 1-2 h at 4 °C. The blocked beads were then incubated with protein lysates for 2h at 4 °C on a rotator. After incubation, beads were washed three times with GST buffer. The bound proteins were eluted by resuspending the beads in 2×sample buffer. For input controls, 20 µg of total protein lysate was mixed with 2× sample buffer. All samples were boiled for 5 min at 95 °C and separated by SDS–PAGE. Proteins were transferred onto polyvinylidene difluoride (PVDF) membranes (EMD Millipore) at 100 V, 350 mA, or 50 W for 90 min. Membranes were blocked overnight at 4 °C in TBST (20 mM Tris-HCl, 150 mM NaCl, 0.1% (v/v) Tween-20, pH 7.6) containing 5% (w/v) non-fat dry milk. Blots were incubated with primary antibodies diluted in TBST containing 5% milk for 1 h at room temperature or overnight at 4 °C. After primary incubation, membranes were sequentially washed for 10 min each in TBST, TBST supplemented with 0.5 M NaCl, and TBST containing 0.5% Triton X-100. Membranes were then incubated with horseradish peroxidase (HRP)-conjugated secondary antibodies diluted in TBST with 5% milk for 1 h at room temperature, followed by three final washes in TBST. Signal detection was performed using an enhanced chemiluminescence (ECL) kit (Bio-Rad Laboratories) and visualized with a Bio-Rad imaging system.

Analysis of Western blot signals was performed using Image Lab software (Bio-Rad). For each blot, the longest exposure without signal saturation was used for quantification. In pull-down experiments, band intensities of experimental samples were normalized to those of the corresponding directly pulled-down protein (e.g., EGFP-tagged proteins in GFP pull-downs). For analyses comparing tagged constructs, signal intensities were first normalized to the corresponding input bands to account for differences in expression levels, followed by normalization to the pull-down control. Normalized values were expressed relative to the control condition, which was set to 100% across all replicates.

### Recombinant proteins purification and interaction assay

The plasmids encoding the GST- or TwinStrep-tagged proteins were transformed into Cheminally competent BL21-CodonPlus(DE3)-RP (Agilent) E. coli cells. These bacteria were cultured in 1L LB broth with 2% D-glucose and 100µg/ml ampicillin and incubated at 37°C. Once the OD_600_ reached 0.4-0.6, the protein expressions were induced with 1mM IPTG and incubated overnight in an 18°C shaker. Once the OD_600_ reaches 1.2, the bacterium cultures were centrifuged for 10 mins at 8000 xg at 4°C. The pellets were resuspended in 10ml PBS (for GST-tagged proteins) or buffer W (IBA Lifesciences; for TwinStrep-tagged proteins) supplemented with protease inhibitor cocktail mini tablet without EDTA (Roche). The bacterial suspensions were then lysed using a JENCONS sonicator at 15W for 3 minutes on ice. Lysates were centrifuged at 39,000 x g for 20 mins to remove the cellular debris. The GST and TwinStrep fusion proteins were purified using glutathione-Sepharose (GE Healthcare) or Strep-Tactin®XT (IBA Lifescience). Purified TwinStrep-tagged proteins were eluted using Buffer BXT (IBA Lifescience). To confirmed successful protein purification, samples were separated by SDS-PAGE and stained using Rapid Blue Colloidal Coomassie Stain (Severn Biotech) following manufacturer’s instructions.

To examine interactions between purified recombinant proteins, glutathione bead attached GST-tagged proteins (GST-empty or GST-LRP5/6) were first washed with 1 ml kinase reaction buffer (KRB; 25 mM Tris-HCl, pH 7.5, 10 mM MgCl₂, 1 mM DTT) and centrifuged at 2,500 × g for 2 min. Beads were dephosphorylated with 10 µl Quick CIP at 37 °C for 20 min, followed by blocking in KRB containing 1% BSA at 4 °C for 1 h with gentle rotation. After washing once with 1 ml KRB, beads were incubated with 2 µg of kinase (GSK3β, CK1ε, or both) in KRB supplemented with 0.2 mM ATP at 30 °C for 1 h. The phosphorylated beads were then washed three times with 1 ml KRB (2,500 × g, 2 min each) and subsequently incubated with the target protein at 4 °C for 1h. Following incubation, beads were washed three additional times with KRB and resuspended in 20 µl of 2× sample buffer. Samples were boiled at 95 °C for 5 min and analyzed by SDS–PAGE

### Super TOP Flash (STF) Reporter Assay

Canonical Wnt signalling activity was assessed using the Super TOP Flash reporter assay utilizing the Dual-Glo® Luciferase Reporter System (Promega). Cells were seeded in six-well plates and co-transfected the following day with 400 ng of SuperTOPFlash or SuperFOPFlash firefly luciferase reporter plasmid together with 80 ng of Renilla luciferase plasmid per well. After 24 h incubation at 37 °C, cells were stimulated with either Wnt3a-conditioned medium or control medium for the indicated duration.

Following stimulation, cells were lysed in 1× Passive Lysis Buffer (Promega). Firefly and Renilla luciferase activities were sequentially measured using a luminometer (Promega) with automatic injection of 20 µl of Luciferase Assay Reagent II (LAR II) and Stop & Glo® Reagent (Promega), respectively. Luminescence was recorded with a 1 s delay and a 10 s integration time for each well. Firefly luciferase activity was normalized to Renilla luciferase to control for transfection efficiency. Data represent the mean of at least three independent experiments: each performed in technical triplicate.

### Microscopy

HeLa cells were seeded at a density of 1×10⁵ cells per well and incubated for 24 hours at 37°C with 5% CO₂. Plasmid transfections were performed using the X-tremeGENE 360 transfection reagent, as described in the transfection section. The next day, 35mm μ-dishes (Ibidi 81158) were coated with 10μg/ml fibronectin (Merck Life Science UK; F1141) in PBS for one hour at room temperature followed by a PBS wash. The transfected HeLa cells were plated onto these dishes in phenol red-free DMEM at a density of 1×10⁵ cells per well. The cells were incubated at 37°C with 5% CO₂ and allowed to spread for 3-4 hours. Imaging was performed using an inverted IX81 widefield microscope (Olympus) equipped with an environmental chamber (37°C and 5% CO_2_), controlled by MetaMorph software. Images were captured at 100x magnification. Image processing was conducted using FIJI.

### LRP6 endocytosis assay

To quantify LRP6 endocytosis, hTERT-RPE1 cells were seeded in 10-cm tissue culture dishes at a density of 1 × 10⁵ cells and cultured for 24 h at 37 °C in 5% CO₂. Cells were transfected using polyethyleneimine (PEI) and incubated for an additional 24 h. Transfected cells were then re-seeded into new 10-cm dishes at 2.35 × 10⁶ cells per dish and cultured overnight.

For surface biotinylation, cells were washed twice with ice-cold PBS and incubated with 5 ml of cold PBS containing 0.2 mg/ml Sulfo-NHS-Biotin for 30 min at 4 °C with gentle agitation. After incubation, cells were washed twice with cold PBS to remove excess reagent. Where indicated, cells were stimulated with the appropriate conditioned medium or purified Wnt3a (5036-WN-010, R&D Systems) for the desired duration. For samples used to assess internalized (rather than total surface) protein, surface biotin was removed by incubating cells with 50 ml of reduction buffer (50mM Glutathione, 75mM NaCl, 10mM EDTA, 1% BSA and 75mM NaOH) for 30 min at 4 °C. Residual reactive groups were quenched by two successive 5 min incubations in iodoacetamide (IAA) buffer (5mg/ml Iodoacetamide) at 4 °C.

Following the final wash, cells were lysed in 500 µl of GST lysis buffer and clarified by centrifugation at 17,000 × g for 15 min at 4 °C. To capture biotinylated proteins, 96-well Maxisorp plates (Thermo Fisher Scientific) were pre-coated with 0.5 µg/ml anti-LRP6 antibody (100 µl per well) in buffer 1 (0.05M Na_2_CO_3_ pH9.6) overnight at 4 °C and blocked for 1 h at room temperature with PBS containing 0.1% Tween-20 and 5% BSA. Clarified cell lysates were added to the antibody-coated wells and incubated overnight at 4 °C.

The next day, plates were washed four times with PBS containing 0.1% Tween-20 and incubated with streptavidin–HRP conjugate for 1 h at room temperature. After four additional washes, bound biotinylated proteins were detected by adding 200 µl of detection buffer (0.8 mg/ml o-phenylenediamine dihydrochloride and 0.009% H₂O₂). Absorbance at 450 nm was recorded using an ELISA plate reader every 5 min for a total of 75 min.

<u>Xenopus experiments:</u> *Xenopus laevis* embryos were obtained using standard methods (*80*, *81*). Synthetic mRNA was generated from cDNA constructs carrying *Wnt8D, FP4-mito* and *AP4-mito* cDNAs described above, using mMESSAGE mMACHINE SP6 transcription kits (Thermo Fisher). Microinjections were performed at the 2-cell stage into the animal hemisphere. Embryos were then cultured to stage 30 or stage 40 as indicated and scored according to Kao & Elinson dorsoanterior index (DAI). Experiments were performed three times with embryos obtained from at least two females each time.

### Statistical analysis

Statistical analyses were performed using GraphPad Prism software. Data are presented as mean ± SEM. For normally distributed data, *p* values were calculated using two-way ANOVA, one-way ANOVA with Dunnett’s post hoc test, or Student’s *t*-test for multiple and single comparisons, respectively. The respective statistical tests are disclosed in figure legends.

**Fig. S1.**
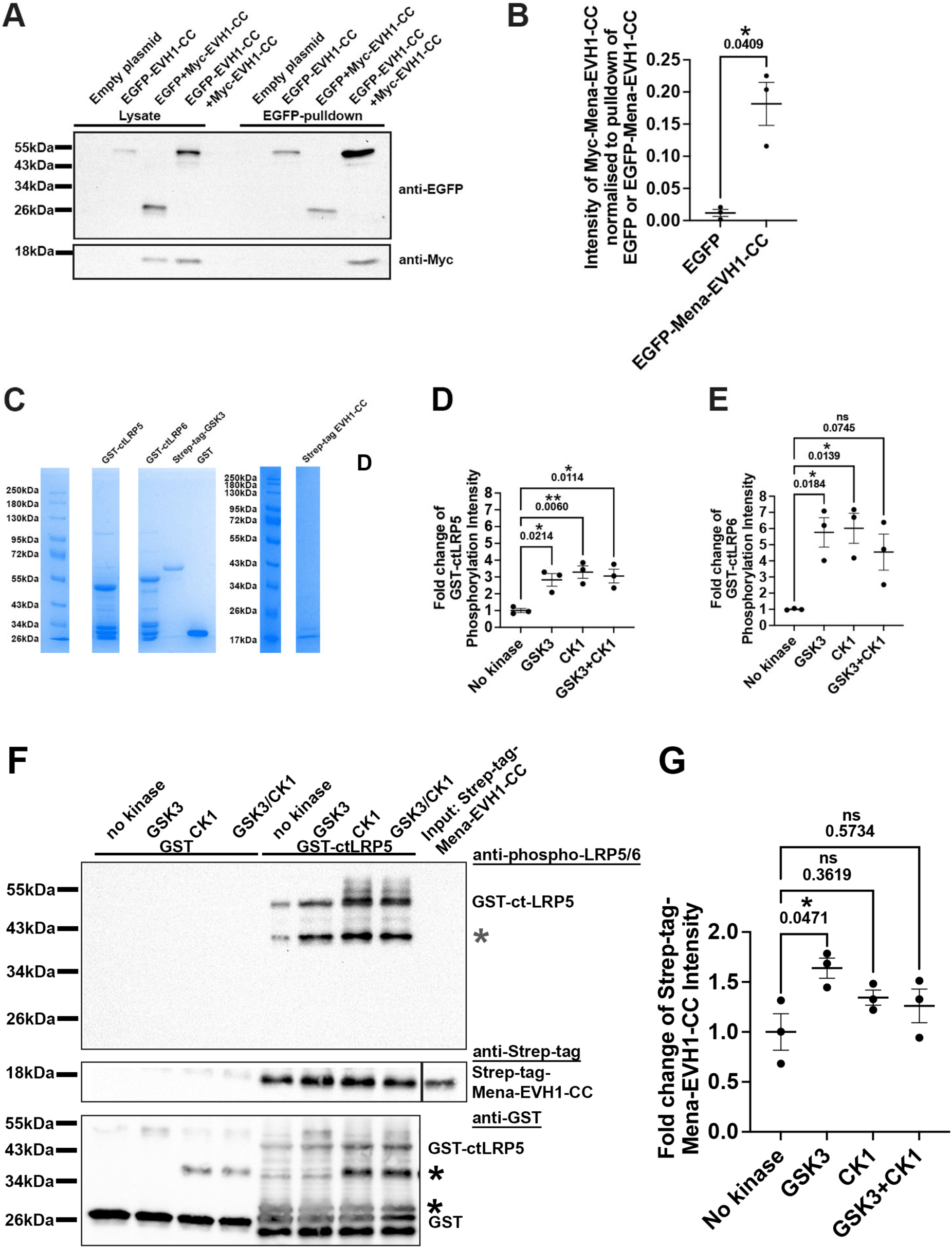
Validation of oligomerization of Mena-EVH1-CC construct and GST-ctLRP5/6 purification and phosphorylation. **(A)** HEK293FT cells were transfected with empty vector, EGFP, EGFP with Myc-Mena-EVH1-CC and EGFP-Mena-EVH1-CC with Myc-Mena-EVH1-CC. EGFP nanobody agarose was used for pulldowns from cell lysates and analysed by western blotting using anti-Myc and anti-EGFP antibodies. Shown is one representative blot from a total of 3 independent repeats. **(B)** Quantification of Myc-Mena-EVH1-CC band intensity of blot (B) normalized to the corresponding EGFP-Mena-EVH1-CC or EGFP band intensity. Data is plotted as mean ± SEM. Paired t-test. **(C)** Coomassie blue stained SDS-PAGE gel depicting the purified GST-ctLRP5, GST-ctLRP6, Strep-tag-GSK3, GST, (left from the same gel) and Strep-tag-EVH1-CC (right from another gel). **(D,E)** Quantification of phospho-ctLRP5 (D) and phospho-ctLRP6 band intensity from Fig. 1 (A, top panel) and (C, top panel). One-way ANOVA, Tukey’s, from 3 independent repeats. **(F,G)** Immobilised, purified GST-tagged LRP5 or GST only as negative control was in vitro phosphorylated with either GSK3, CK1, or both and incubated with purified Mena Strep-tag-EVH1-CC. Phosphorylation was assessed using an anti-phospho-LRP5 antibody (top panel). Interaction was visualized in a western blot against the strep-tag (middle panel) or GST (lower panel). Note some degradation of GST-ctLRP6 occurs due to its unstructured nature indicated in (F) as *. **(G)** Quantification of Strep-tag-Mena-EVH1-CC band intensity from (F). One-way ANOVA, Tukey’s; Mean ± SEM; three independent biological repeats.

**Fig. S2.**
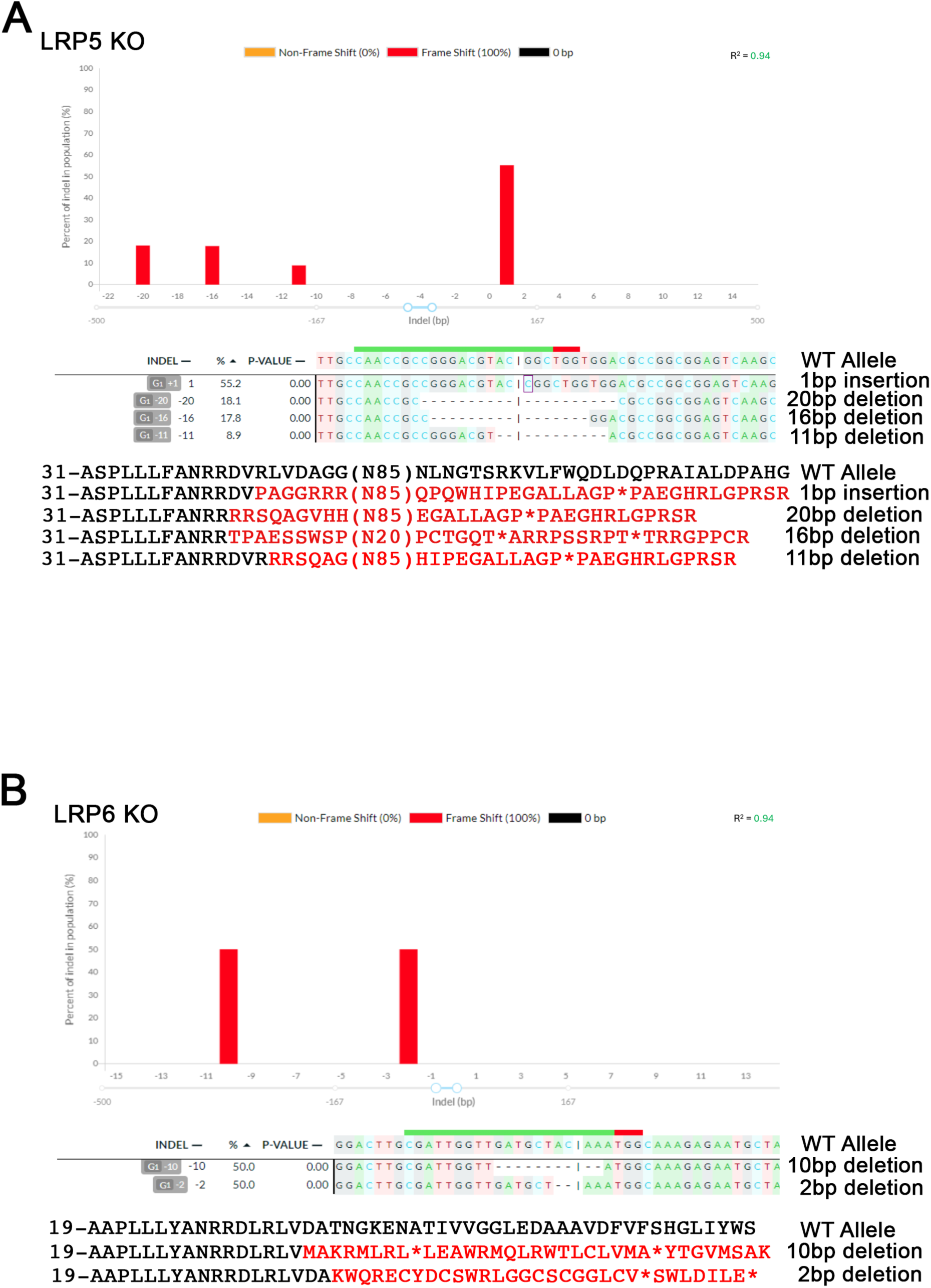
Validation of the LRP5 and LRP6 CRISPR knockout HEK193 cell lines. **(A – B)** Genomic DNA was isolated from wild-type HEK293 cells as well as potential LRP5 (A) or LRP6 (B) KO clones, and the region of DNA encompassing the expected cut site in exon 2 was amplified by PCR. These amplicons were sequenced, and DNA Sequences of knock-out clones were compared to WT sequence using the DECODR web tool. (B) 100% of alleles in the LRP5 KO clone cause a frame shift with deletions of 1, 11, 16, 20 bp resulting frame shifts and a premature termination codon. (B) 100% of alleles in the LRP6 KO clone cause a frame shift with deletions of 2 and 10 bp. Both mutations result in a frame shift and a premature termination codon. For both panels the amino acid sequence of exon 2 around the cut site from aa-31 (NP_002326.2) for LRP5 and from aa-19 (NP_001401173.1) for LRP6 is shown in the wild-type and mutant alleles. Altered amino acid sequence as a result of the frame shift is shown in red and the premature termination codon is represented with a ‘*’.

**Fig. S3.**
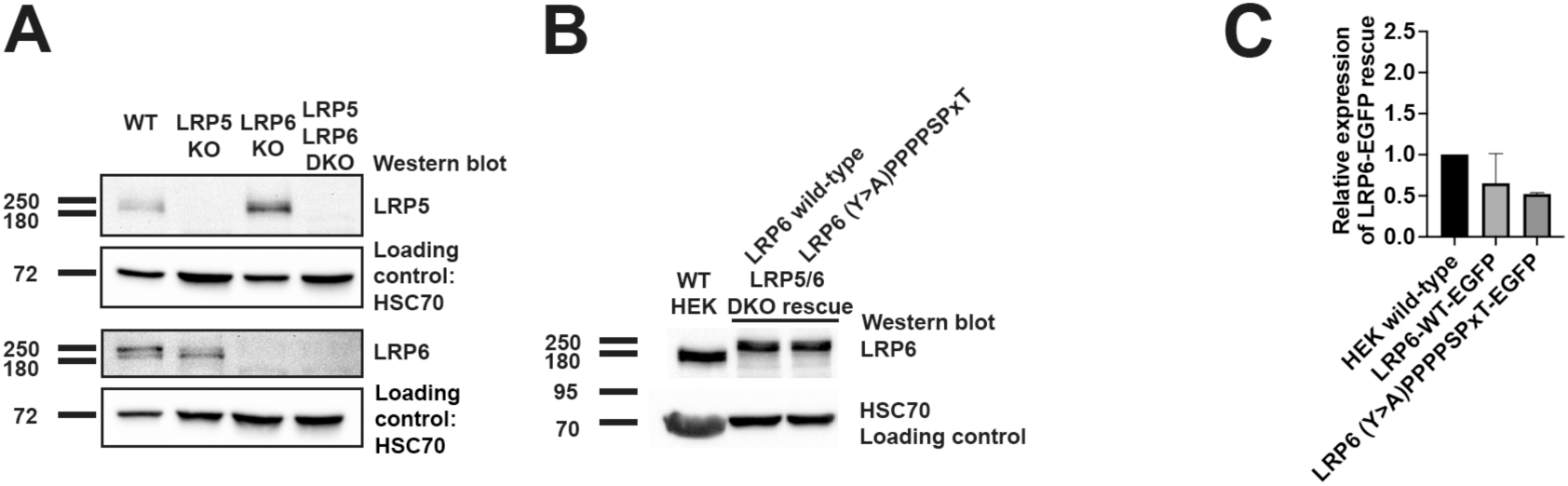
Validation of LRP5 and LRP6 double knockout HEK293 cell line and its rescue with wild-type or mutated LRP6. **(A)** LRP5 or LRP6 single and LRP5/6 double knockout HEK293 cells were lysed and expression of endogenous LRP5 or LRP6 was tested in a western blot. HSC70 served as the loading control. **(B)** LRP5/6 double knockout HEK293 cells were transfected with either wild-type or LRP6 cDNA mutated in the Ena/VASP EVH1 domain binding sites. Expression levels were compared to wild-type, non-transfected HEK293 cells in a western blot for endogenous LRP6. HSC70 served as the loading control. **(C)** Quantification of the western blot from (B).

**Fig. S4.**
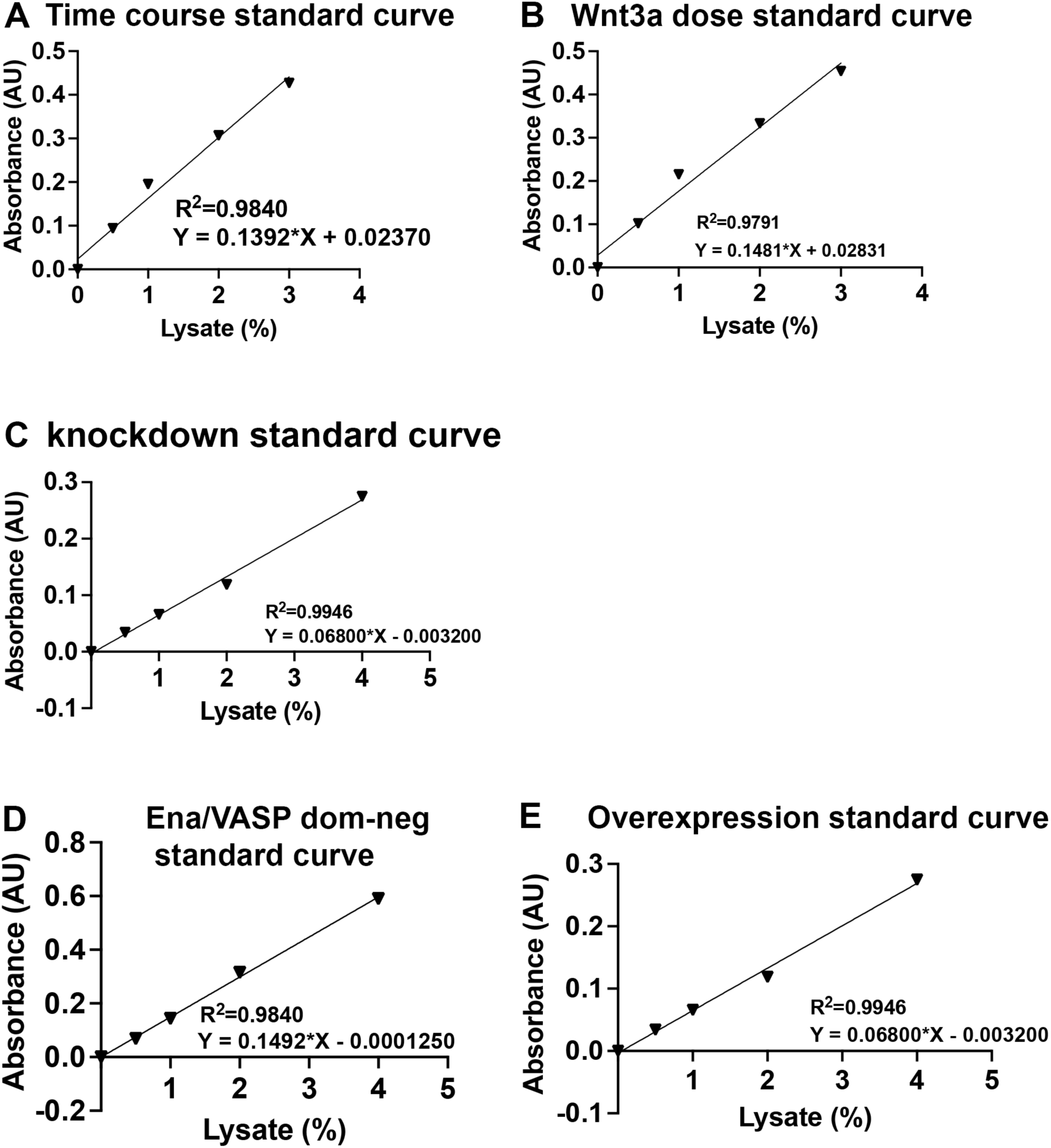
Standard curves for the ELISA quantification of endogenous LRP6 endocytosis. **(A-E)** Linear regression standard curves for the quantification of endogenous LRP6 endocytosis using hTERT-RPE1 cells from Figure 3 for (A) time course of control or Wnt3a conditioned media (CM) (relates to Fig. 3C) (B) experiment to determine lowest concentrations of purified Wnt3a needed to induce LRP6 endocytosis (relates to Fig. 3D) (C) knockdown of endocytosis regulators (relates to Fig. 3E) (D) Ena/VASP dominant negative experiment (relates to Fig. 3F) (E) EGFP-tagged Mena, VASP, EVL overexpression experiment (relates to Fig. 3G).

**Fig. S5.**
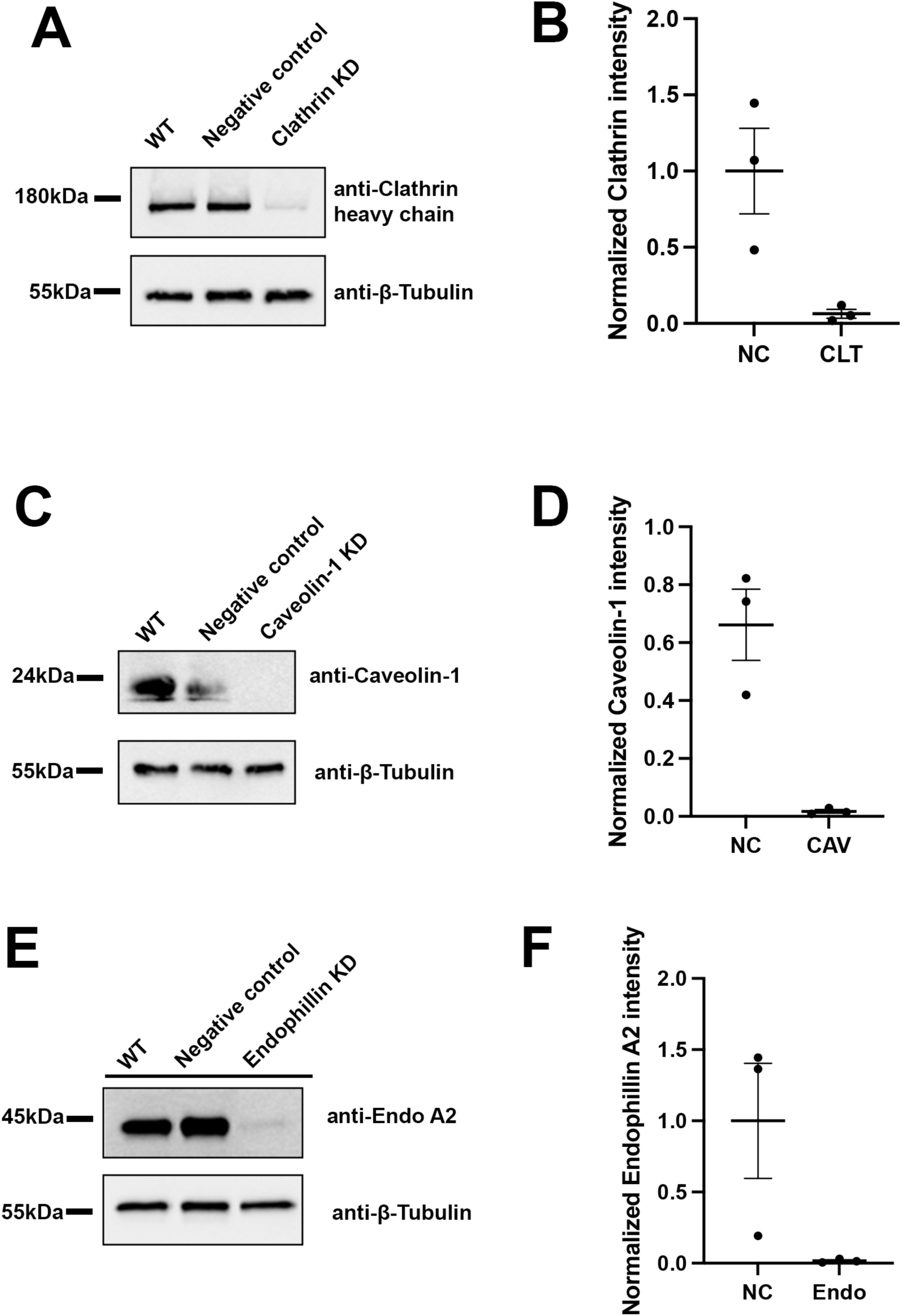
Evaluation of knockdown efficiency of endocytosis mediators. **(A-F)** hTERT-RPE1 cells were transfected with siRNAs specific to (A,B) clathrin heavy chain, (C,D) Caveolin-1, (E,F) Endophilin A1-3. Cell lysates were probed in a western blot with respective antibodies and reprobed with anti-beta-tubulin antibodies as a loading control. (B,D,F) Quantification of loading control normalized band intensity from (A) clathrin heavy chain, (C) Caveolin-1, (E) Endophilin A2. (relates to Fig. 3E).

**Table S1.**
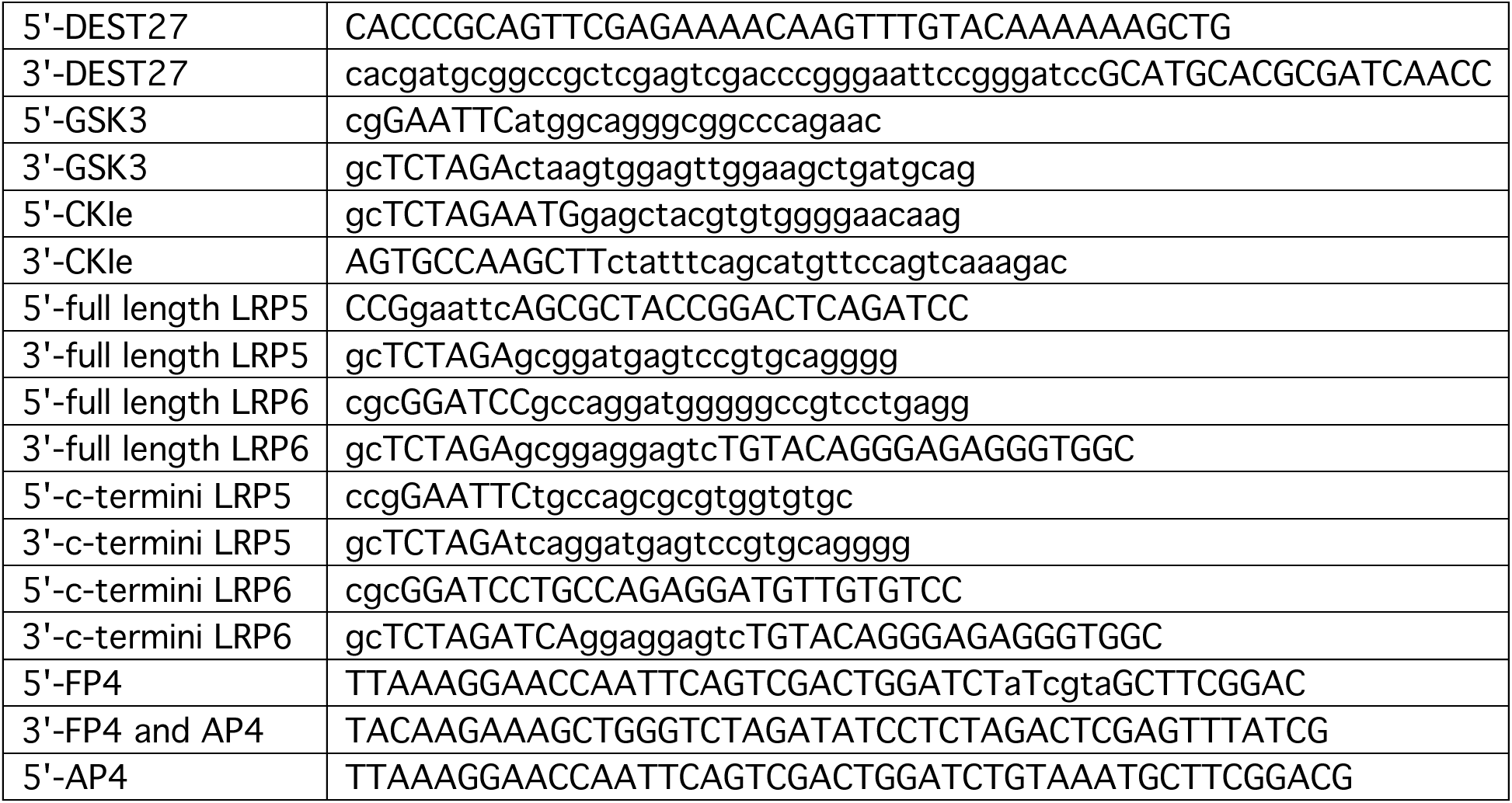
Primer sequences used for cloning.

**Table S2.**
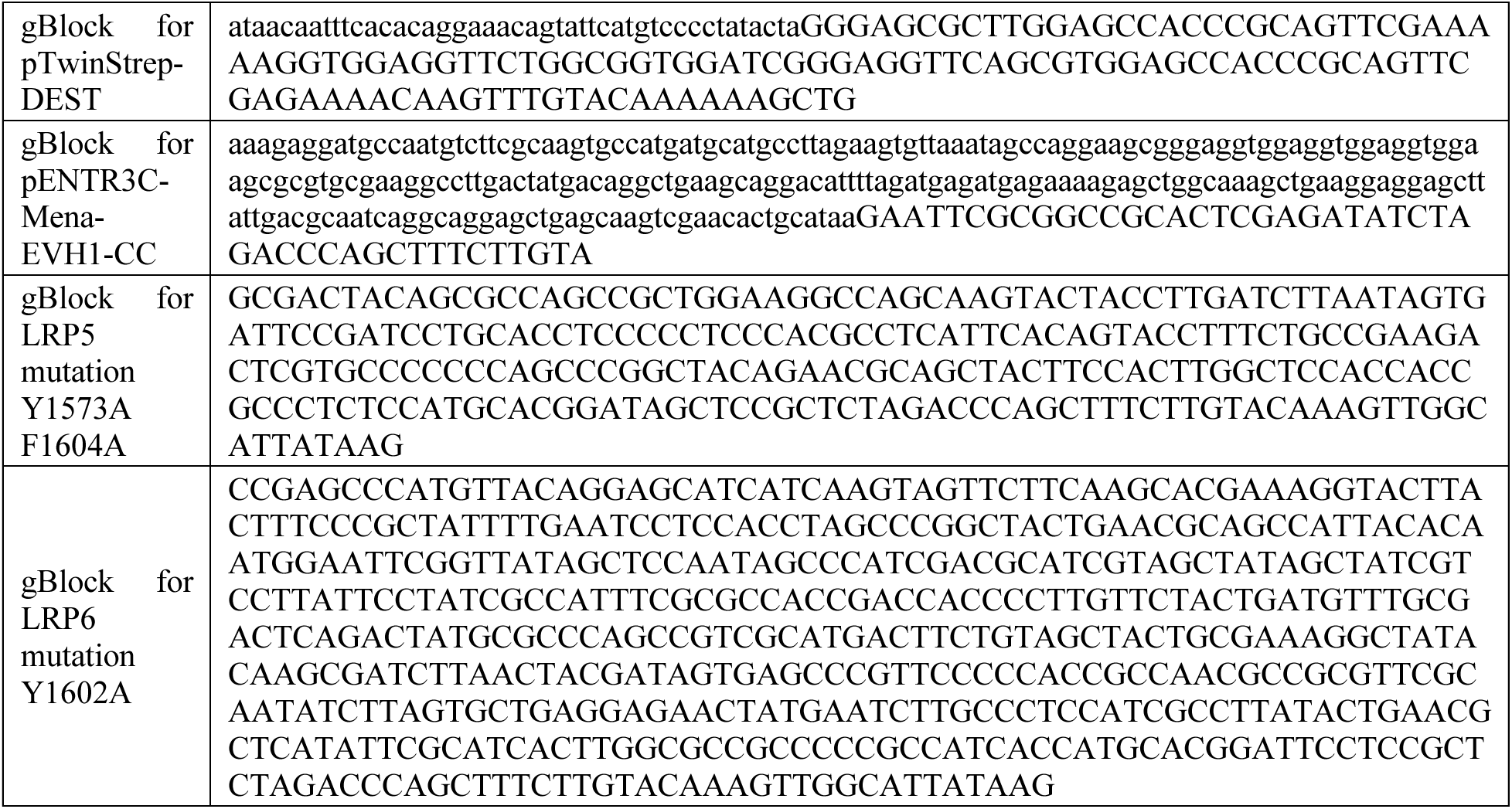
gBLOCK synthesized DNA fragments. Type or paste caption here. Create a page break and paste in the table above the caption.

